# Testing theoretical minimal genomes using whole-cell models

**DOI:** 10.1101/2020.03.26.010363

**Authors:** Joshua Rees-Garbutt, Jake Rightmyer, Oliver Chalkley, Lucia Marucci, Claire Grierson

**Affiliations:** BrisSynBio, University of Bristol, Bristol BS8 1TQ, UK; School of Biological Sciences, University of Bristol, Bristol Life Sciences Building, 24 Tyndall Avenue, Bristol, BS8 1TQ, UK; Bristol Centre for Complexity Sciences, Department of Engineering Mathematics, University of Bristol, Bristol, BS8 1UB, UK; Department of Engineering Mathematics, University of Bristol, Bristol BS8 1UB, UK; School of Cellular and Molecular Medicine, University of Bristol, Bristol BS8 1 UB, UK

## Abstract

The minimal gene set for life has often been theorised, with at least ten produced for *Mycoplasma genitalium* (*M. genitalium*). Due to the difficulty of using *M. genitalium* in the lab, combined with its long replication time of 12 - 15 hours, none of these theoretical minimal genomes have been tested, even with modern techniques. The publication of the *M. genitalium* whole-cell model provided the first opportunity to test them, simulating the genome edits *in-silico*. We simulated eight minimal gene sets from the literature, finding that they produced *in-silico* cells that did not divide. Using knowledge from previous research, we reintroduced specific essential and low essential genes *in-silico*; enabling cellular division. This reinforces the need to identify species-specific low essential genes and their interactions. Any genome designs created using the currently incomplete and fragmented gene essentiality information, will very likely require *in-vivo* reintroductions to correct issues and produce dividing cells.

## Introduction

Genome engineering builds on historical gene essentiality research. The sequencing of small bacterial genomes (Fleischmann et al., 1995; Fraser et al., 1995) led to comparative genomics; then, as genome sequences completed increased, minimal gene sets (Forster & Church, 2006; R. Gil et al., 2004; Rosario Gil, 2015; J. I. Glass et al., 2006; Huang et al., 2013; Hutchison et al., 1999; Karr et al., 2012; Mushegian & Koonin, 1996; Shuler et al., 2012; Tomita et al., 1999) were hypothesised. Minimal genomes are reduced genomes where no single gene can be removed without loss of reproduction (John I. Glass et al., 2017), given an appropriately rich medium and no external stresses, and focusing solely on protein-coding genes. For a recent review of gene essentiality, see Rancati *et al*. (Rancati et al., 2018).

*M. genitalium* is the focal point of minimal gene set creation due its naturally small genome size (0.58mb and 525 genes) and early sequenced genome (Fraser et al., 1995). Minimal gene sets are designed using three different approaches: protocells, comparative genomics, and single gene knockouts. Protocell designs (Dzieciol & Mann, 2012) are not expected to function as true biological cells, instead functioning as a self-replicating, membrane-encapsulated collection of biomolecules (Forster & Church, 2006). Comparative genomics (Koonin, 2003) compares multiple species to identify common genes. This is complicated (John I. Glass et al., 2017; Lagesen et al., 2010) by non-orthologous gene displacements, i.e. independently evolved or diverged proteins that perform the same function but are not recognisably related (John I. Glass et al., 2017; Mushegian & Koonin, 1996), which can result in the removal of a large number of genes essential to one species. Design using single gene essentiality classifications should, in theory, not remove any essential genes; but if transposon mutagenesis is used, variance from different transposon variants, antibiotic resistance genes, and growth periods can result in differing essentiality classifications (J. I. Glass et al., 2006; Juhas et al., 2011).

Ten minimal gene sets were found in the literature that were designed with *M. genitalium* genes (Forster & Church, 2006; R. Gil et al., 2004; Rosario Gil, 2015; J. I. Glass et al., 2006; Huang et al., 2013; Hutchison et al., 1999; Karr et al., 2012; Mushegian & Koonin, 1996; Shuler et al., 2012; Tomita et al., 1999). Due to the difficulty of using *M. genitalium* in the lab (Reich, 2000), combined with its long replication time of 12 - 15 hours (Benders et al., 2010; John I. Glass et al., 2017; Hutchison et al., 2016), none of these theoretical minimal genomes have been tested, even with modern techniques (Benders et al., 2010). The publication of the *M. genitalium* whole-cell model (Karr et al., 2012) provided the first opportunity to test them, simulating the genome edits *in-silico*.

The *Mycoplasma genitalium* whole-cell model (Karr et al., 2012) was the first existing model of a cell’s individual molecules that includes the function of every known gene product (401 of the 525 *M.genitalium* genes), making it capable of modelling genes in their genomic context. Previously, it has been used to investigate discrepancies between the model and real-world measurements (Karr et al., 2012; Sanghvi et al., 2013), design synthetic genetic circuits in the context of the cell (Purcell et al., 2013), make predictions about the use of existing antibiotics against new targets (Kazakiewicz et al., 2015), and produce *in-silico* minimal genomes (Rees-Garbutt et al., 2020).

Of the ten sets, two of the sets (Gil *et al*. (2004)(R. Gil et al., 2004)and Shuler *et al.(Shuler et al., 2012))* were excluded as they were considered derivative of the Gil (2014) set (Rosario Gil, 2015); four genes differ in the Shuler *et al*. set (MG_056, MG_146, MG_388, MG_391) and four genes are absent in the Gil *et al*. (2004) set (MG_009, MG_091, MG_132, MG_460). Of the other eight sets, two (Tomita *et al*. (Tomita et al., 1999) and Church *et al*. (Forster & Church, 2006)) were designed as protocells, three (Mushegian and Koonin (Mushegian & Koonin, 1996), Huang *et al*. (Huang et al., 2013), and Gil (R. Gil et al., 2004; Rosario Gil, 2015)) from comparative genomics, and three (Hutchison *et al*. (Hutchison et al., 1999), Glass *et al*. (J. I. Glass et al., 2006), and Karr *et al*. (Karr et al., 2012)) from single gene essentiality experiments.

We simulated eight minimal gene sets from the literature (Forster & Church, 2006; Rosario Gil, 2015; J. I. Glass et al., 2006; Huang et al., 2013; Hutchison et al., 1999; Karr et al., 2012; Mushegian & Koonin, 1996; Tomita et al., 1999), testing them for the first time. This was to assess their functionality as extant and untested genome designs, but also to produce new data for designing genomes to evaluate the differing approaches of focusing on species-specific data or focusing on identifying core genes and common functions across species.

## Results

We adapted eight minimal gene sets for simulation within the *M. genitalium* whole-cell model. To increase clarity, we renamed the sets after the main location where the set was constructed (Table 1). The Bethesda set directly compared *M. genitalium* and *Haemophilus influenzae* genomes (gram-positive and gram-negative bacteria, respectively) (Mushegian & Koonin, 1996). The Rockville set applied global transposon mutagenesis to *M. genitalium in-vivo* to identify non-essential genes (Hutchison et al., 1999). The Fujisawa set constructed an *in-silico* hypothetical cell from 127 *M. genitalium* genes using the E-Cell software (Tomita et al., 1999). The Rockville 2 set reapplied global transposon mutagenesis *in-vivo*, isolating and characterising pure clonal populations (J. I. Glass et al., 2006). The Nashville set listed 151 *E.coli* genes (compared to *M. genitalium* genes within the paper) theorised to produce a chemical system capable of replication and evolution (Forster & Church, 2006). The Stanford set was the result of *in-silico* single gene knockouts using the *M. genitalium* whole-cell model (Karr et al., 2012). The Guelph set compared 186 bacterial genomes (Huang et al., 2013), whereas the Valencia set compared *M. genitalium* with genetic data of five insect endosymbionts (Rosario Gil, 2015). The Nashville, Fujisawa, and Stanford sets were unchanged, but the others had between 6 and 44 genes removed (Table 1) either because the genes were unmodelled (the genes’ function were unknown (Karr et al., 2012)) or genes had duplicate entries.

**Table 1.**
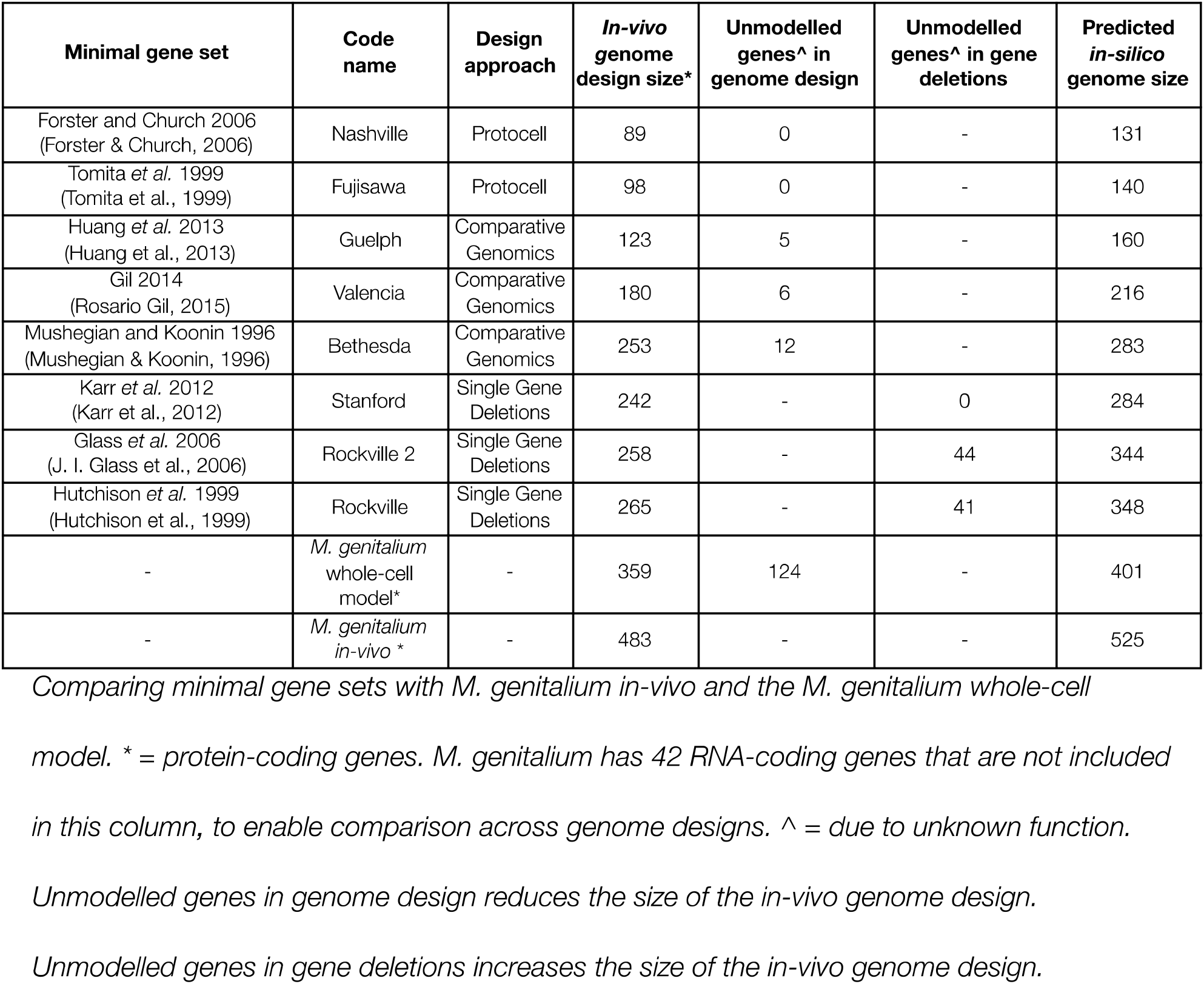
Minimal Gene Sets from the literature.

| Minimal gene set | Code name | Design approach | *In-vivo* genome design size* | Unmodelled genes^ in genome design | Unmodelled genes^ in gene deletions | Predicted *in-silico* genome size |
| --- | --- | --- | --- | --- | --- | --- |
| Forster and Church 2006 (Forster & Church, 2006) | Nashville | Protocell | 89 | 0 | - | 131 |
| Tomita *et al.* 1999 (Tomita et al., 1999) | Fujisawa | Protocell | 98 | 0 | - | 140 |
| Huang *et al.* 2013 (Huang et al., 2013) | Guelph | Comparative Genomics | 123 | 5 | - | 160 |
| Gil 2014 (Rosario Gil, 2015) | Valencia | Comparative Genomics | 180 | 6 | - | 216 |
| Mushegian and Koonin 1996 (Mushegian & Koonin, 1996) | Bethesda | Comparative Genomics | 253 | 12 | - | 283 |
| Karr *et al.* 2012 (Karr et al., 2012) | Stanford | Single Gene Deletions | 242 | - | 0 | 284 |
| Glass *et al.* 2006 (J. I. Glass et al., 2006) | Rockville 2 | Single Gene Deletions | 258 | - | 44 | 344 |
| Hutchison *et al.* 1999 (Hutchison et al., 1999) | Rockville | Single Gene Deletions | 265 | - | 41 | 348 |
| - | *M. genitalium* whole-cell model* | - | 359 | 124 | - | 401 |
| - | *M. genitalium in-vivo* * | - | 483 | - | - | 525 |

*Comparing minimal gene sets with M. genitalium in-vivo and the M. genitalium whole-cell* *model. \* = protein-coding genes. M. genitalium has 42 RNA-coding genes that are not included* *in this column, to enable comparison across genome designs. ^ = due to unknown function.*
*Unmodelled genes in genome design reduces the size of the in-vivo genome design.*
*Unmodelled genes in gene deletions increases the size of the in-vivo genome design.*

The protocell designs (Nashville, Fujisawa) predicted the smallest *in-silico* genome. Guelph contained substantially fewer genes than Valencia and Bethesda, due to comparing 186 species (Huang et al., 2013). Stanford, Rockville, Rockville 2 had similar numbers *of in-vivo* deletions, but Rockville and Rockville 2 had the highest numbers of unmodelled genes (as genes can be disrupted *in-vivo* without knowing the gene’s function). The *in-silico* genomes (and associated gene deletions) of the minimal gene sets are listed in Supplementary Data 1 and 2.

Prior to simulations, we analysed the sets produced by comparative genomics and single gene essentiality and found 96 genes they had in common (Supplementary Data 1 (col L) and 3). We excluded the protocell designs from this comparison due to their much reduced size. 87 of the shared genes were classified as essential (genes that when removed *in-silico* stopped the cell from dividing, classified *in-silico* previously (Karr et al., 2012; Rees-Garbutt et al., 2020)), eight were non-essential (removal did not prevent successful cell division), and one gene was unmodelled.

The 87 essential genes affect a range of cellular functions including: DNA (repair, supercoiling, chromosome replication, nucleotide synthesis/modification, sigma factors, ligation, transcription termination, and DNA polymerase); RNA (ribosome proteins, translation initiation factors, tRNA modification, ribonucleases, and RNA polymerase); and cellular processes (protein folding/modification, protein shuttling, protein membrane transport, metabolic substrates production/recycling, redox signalling, oxidation stress response, and the pyruvate dehydrogenase complex). Of the eight non-essential genes, four (MG_048, MG_072, MG_170, MG_297) are associated with the SecYEG complex (du Plessis et al., 2011) (protein transport across or into the cell membrane), while MG_172 removes protein synthesis targets from synthesised proteins, MG_305 and MG_392 assist in late protein folding, and MG_425 processes ribosomal RNA precursors. Although these eight genes are singly non-essential (by single gene deletion *in-silico* (Karr et al., 2012) and *in-vivo* (J. I. Glass et al., 2006)) they all play a part in essential functions, hence their inclusion.

We also identified 14 genes deleted by all eight minimal gene sets (Supplementary Data 2 (col L) and 4). The functions of these genes include: fructose import, host immune response activation, chromosomal partition, amino acid transport, antibody binding, phosphonate transport, external DNA uptake, DNA repair, rRNA modification, membrane breakdown, toxin transport, quorum sensing, and a restriction enzyme. These had been previously classified as non-essential by single gene deletion *in-silico* (Karr et al., 2012) and *in-vivo* (J. I. Glass et al., 2006). We placed these 14 common genes in an Agreed set’ and a genome with these genes removed was also simulated.

We simulated each minimal gene set in the *M. genitalium* whole-cell model and found that every set, including the Agreed set, produced a non-dividing *in-silico* cell (30 *in-silico* replicates, Supplementary Data 5).

Analysis found that every one of the sets deleted essential genes (classified *in-silico* previously (Karr et al., 2012; Rees-Garbutt et al., 2020)): Nashville deleted 121, Fujisawa deleted 112, Guelph deleted 107, Valencia deleted 69, Bethesda deleted 34, Rockville and Rockville 2 both deleted 9, and Stanford deleted 3 (Supplementary Data 6-14). This is especially surprising for the single gene essentiality minimal gene sets. For the Rockville sets, this is likely due to transposon mutagenesis issues. Rockville labelled six genes as non-essential in 1999, subsequently labelled essential in 2006. Additionally, Rockville grew cells in mixed pools with DNA isolated from these mixtures rather than from isolated pure colonies of cells (J. I. Glass et al., 2006). Whereas for Rockville 2, the engineers of *JCVI-Syn3.0* used the same transposon mutagenesis protocol (Hutchison et al., 2016), but found that they had to improve upon it due to incorrect identification of essentiality. For the Stanford set, the removal of MG_203, MG_250, and MG_470 is likely due to averaging multiple simulation’s data together before computational assessment, genes later found to be essential (Supplementary Table 3(Rees-Garbutt et al., 2020)).

In an attempt to restore *in-silico* division, we reintroduced essential genes to the minimal gene sets (Figure 1). Based on previous research (Rees-Garbutt et al., 2020), we also reintroduced low essential genes (i.e. genes dispensable in some contexts, such as redundant essential genes and gene complexes (Rancati et al., 2018)). We did this by comparing the gene content of the individual minimal gene sets with a complete list of the *M. genitalium in-silico* genes and their essentiality classifications (Rees-Garbutt et al., 2020) (Supplementary Data 6-14). For example, the original Agreed set removed low essential genes MG_291 (phosphonate transport) and MG_412 (phosphate transport); by disrupting both these processes, the *in-silico* cell has no functioning source of phosphate, which has been established previously (Rees-Garbutt et al., 2020). By reintroducing set-specific genes (Table 2, Supplementary Data 15), each modified set, including the Agreed set, was able to produce a dividing cell *in-silico* (30 *in-silico* replicates, Supplementary Data 5).

**Figure 1.**
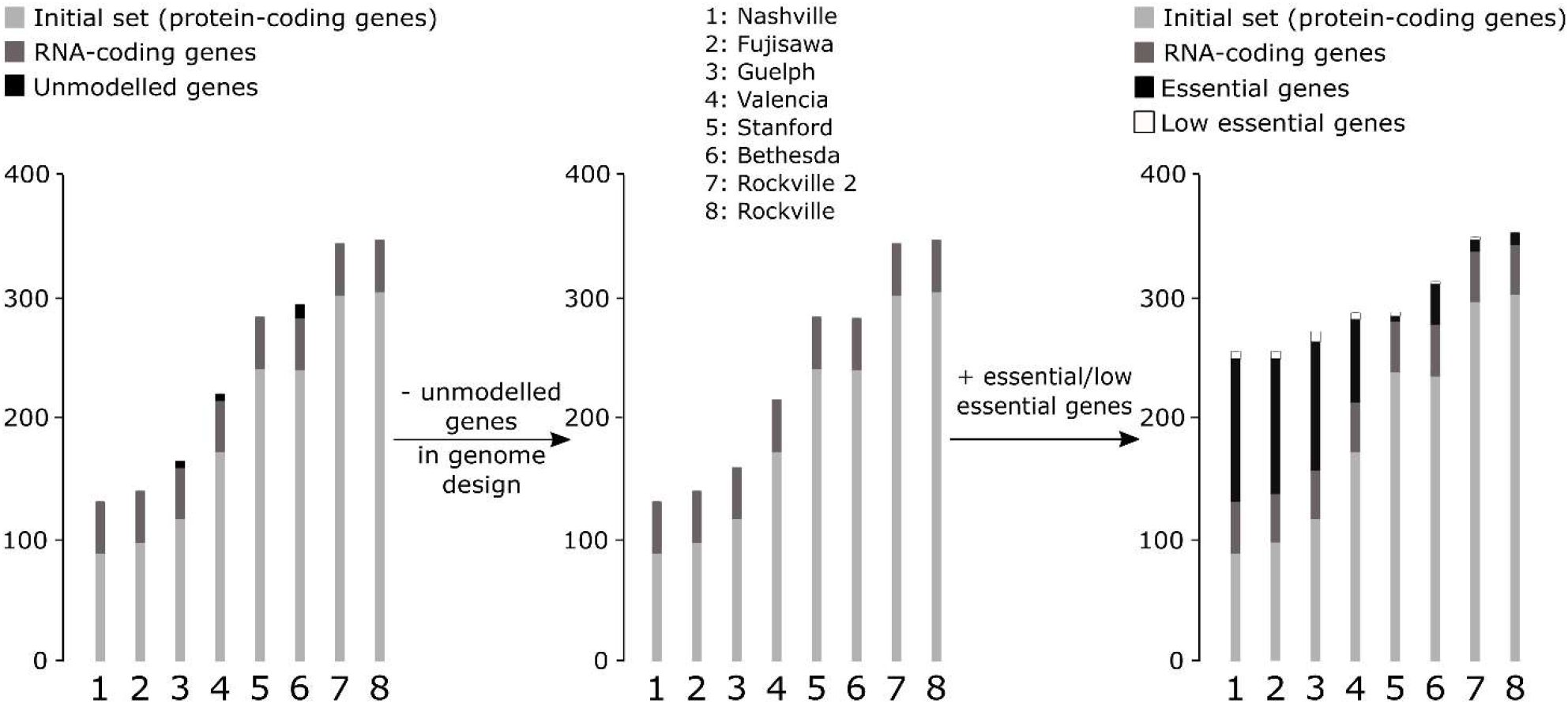
Testing and restoring the minimal gene sets. 124 genes were unmodeled in the M. genitalium whole-cell model due to unknown function. Essential genes are required by the cell to enable survival until successful division. Low essential genes are required by the cell in certain genomic and environmental contexts to enable survival to successful division. The restored minimal gene sets produced dividing in-silico cells. Greater detail on the minimal gene sets is provided in Table 1 and 2.

**Table 2.** Reintroducing genes to produce division. Minimal gene sets before and after the reintroduction of essential and low essential genes dividing in-silico cells. *Size of in-silico genome includes both the 359 protein-coding genes and the 42 RNA-coding genes in the M. genitalium whole-cell model, to provide a better comparison for in-vivo experiments and future work.

| Code name | *In-silico* gene deletions (cell did not divide) | *In-silico* gene deletions (cell divided) | Essential genes reintroduced | Low essential genes reintroduced | Size of *in-silico* genome * |
| --- | --- | --- | --- | --- | --- |
| Nashville | 270 | 142 | 121 | 7 | 259 |
| Fujisawa | 261 | 142 | 112 | 7 | 259 |
| Guelph | 241 | 128 | 107 | 6 | 273 |
| Valencia | 185 | 110 | 69 | 6 | 291 |
| Stanford | 117 | 109 | 3 | 5 | 292 |
| Bethesda | 118 | 82 | 34 | 2 | 319 |
| Rockville 2 | 57 | 45 | 9 | 3 | 356 |
| Rockville | 53 | 43 | 9 | 1 | 358 |
| Agreed | 14 | 13 | 0 | 1 | 388 |

In an attempt to gain further understanding, we investigated what processes the nine repaired minimal gene sets removed using gene ontology (GO) biological process terms, and we compared the repaired minimal gene sets to the *in-silico M. genitalium* minimal genomes we produced previously (Rees-Garbutt et al., 2020). The smallest repaired *in-silico* genomes (Nashville and Fujisawa, 259 genes) were larger than the prior novel *in-silico* minimal genomes (256 and 237 genes), and both removed the low essential genes related to phosphonate transport, relying on inorganic phosphate being transported into the cell (Supplementary Table 6(Rees-Garbutt et al., 2020)). The other sets had different designs that, due to not systematically targeting non-essential genes, resulted in non-essential genes remaining in the genome, making them subsets of the repaired protocell sets and the prior *in-silico* minimal genomes (Supplementary Data 16). As such, the GO terms were also subsets and did not deviate from what we would expect to produce a dividing *in-silico* cell (Supplementary Data 17-25).

Analysis of the repaired sets found that 31 genes were reintroduced into five or more of the minimal gene sets (Supplementary Data 26). 26 were essential and 5 were low essential (Supplementary Table 5(Rees-Garbutt et al., 2020)). The corresponding cellular functions included: DNA (polymerase subunits, thymidine insertion, recycling of pyrimidine, chromosome segregation); RNA (polymerase subunit, tRNA modification, the 5OS and 3OS ribosomal subunits); transporters (cobalt, phosphonate, potassium); production (NAD, flavin, NADP, fatty acid/phospholipids); and dehydrogenation (glycerol and alpha-keto acids). Of the 26 reintroduced essential genes, 19 were already present in the single gene deletion minimal gene sets (Stanford, Rockville, Rockville 2). MG_I37 (mycobacteria cell wall production) and MG_517 (plasma membrane stability) are genes specifically essential for *Mycoplasma* species, which were only identified as essential by the single gene deletion minimal gene sets. A further five genes were involved in cobalt transport, which increases the rates of DNA synthesis, fatty acid metabolism, and amino acid metabolism, and were also not identified as essential by the other design methodologies. Of the five reintroduced low essential genes, Bethesda did not delete four, likely due to the direct comparison of two genomes resulting in low essentiality genes being conserved to a greater degree.

We also looked at reintroductions to the protocell sets, as they could outline additional cellular requirements for the successful future unification of independent protocell systems. The genes reintroduced to the Nashville set repaired functions that had been reduced (translation, glycolytic process, protein folding) and restored functions that had been removed including: cell (division, cycle, transport, redox homeostasis), DNA (topological change, transcription), rRNA processing (including pseudouridine synthesis), protein transport, and cellular processes (carbohydrate metabolic, glycerol metabolic, fatty acid biosynthesis, UMP salvage) (Supplementary Data 27). The glycolytic process had the most change, with 10 of 11 genes being reintroduced, and DNA repair had the least, with only one gene being reintroduced (MG_254), however, this did reintroduce single strand DNA break repair. The genes additionally reintroduced to the Fujisawa set included tRNA processing and protein folding (Supplementary Data 28), with 8 out of 10 DNA replication genes reintroduced.

## Discussion

In conclusion, the repaired protocell minimal gene sets (Nashville and Fujisawa) produced the smallest genomes *in-silico* (Table 2), differing by 6 genes (Supplementary Data 29), but required the most gene reintroductions. The repaired comparative genomics minimal gene sets (Guelph, Valencia, Bethesda) required fewer gene reintroductions, presumably because fewer genomes were compared during their original design. Interestingly, Stanford (a single gene essentiality set) produced a smaller *in-silico* genome than Bethesda (a comparative genomics set), as it did not target unmodelled gene deletions and only required eight genes to be reintroduced.

This research has limitations associated with the use of the *M. genitalium* whole-cell model. Through necessity the *M. genitalium* whole-cell model bases some of its parameters on data from other bacteria^8^ and is only capable of modelling a single generation, missing subgenerational gene expression and subsequent essentiality which affects >50% of the genes in some species (Macklin et al., 2020). Additional uncertainty exists around the unknown impact of the unmodelled genes on *in-vivo* experiments, as stated previously (Rees-Garbutt et al., 2020).

The computational predictions we have produced need to be tested in living cells, but with the advancement of gene synthesis and genome transplantation in other *Mycoplasma* species (Benders et al., 2010; Gibson, Benders, Andrews-Pfannkoch, et al., 2008; Gibson, Benders, Axelrod, et al., 2008; Gibson et al., 2010; Karas et al., 2013; Tsarmpopoulos et al., 2016) this is becoming a more realistic proposition for *Mycoplasma* researchers.

Without the ability to identify species-specific low essential genes, any minimal gene sets designed with the current incomplete and fragmented gene essentiality information will require gene reintroductions to produce dividing cells. New research is required to generate the knowledge required to fill these gaps. Currently, genome-scale engineering has not combined *in silico* and *in vivo* research (Jiang et al., 2013; Karr et al., 2012; Rees-Garbutt et al., 2020). With the recent publication of an *E.coli* whole-cell model (Macklin et al., 2020) and the introduction of a synthetic, recoded *E.coli* genome into a cell (Fredens et al., 2019), this will quickly change. The current nature of species-specific gene essentiality information is, potentially, the only remaining barrier to producing custom genomes.

## Methods

These methods have been published previously [27].

### Experimental Design and Statistics

Each simulation was replicated 30 times. Prior publications have completed five repetitions [8], ten or one hundred repetitions [27], depending on data storage capacity and supercomputer usage. These were the main factors in our choice for the number of replications.

The initial conditions for each simulation are randomised within a natural range for *M. genitalium* cells, introducing variation in the behaviour of simulated cells [8], as previously shown [27]. Statistical tests were not used in this research.

### Code Availability

All code created as part of this paper will be made available on Github under a GNU General Public License v3.0 (gpl-3.0). For more information see choosealicense.com/licenses/lgpl-3.0/.

### Data Availability

The databases used to design the *in-silico* experiments, and compare the results to, includes Karr *et al*. (Karr et al., 2012) and Glass *et al*. (J. I. Glass et al., 2006) Supplementary Tables, and Fraser *et al. M. genitalium* G37 genome (Fraser et al., 1995) interpreted by KEGG (Kanehisa & Goto, 2000) and UniProt (Apweiler et al., 2004) as strain ATCC 33530/NCTC 10195. The output .fig files for all simulations referenced will be made available at the group’s Research Data Repository (data-bris) at the University of Bristol.

### Model Availability

The *M. genitalium* whole-cell model is freely available: github.com/CovertLab/WholeCell. The model requires a single CPU and can be run with 8GB of RAM. We run the *M. genitalium* whole-cell model on Bristol’s supercomputers using MATLAB R2013b, with the model’s standard settings. However, we use our own version of the SimulationRunner.m. MGGRunner.m (github.com/GriersonMaruccil_ab/Analysis_Code_for_Mycoplasma_genitalium_whole-cell_model) is designed for use with supercomputers that start hundreds of simulations simultaneously. It artificially increments the starting time-date value for each simulation, as this value is subsequently used to create the initial conditions of the simulation. Our research copy of the whole-cell model was downloaded 10th January 2017.

### *M. genitalium in-silico* Environmental Conditions

*M. genitalium* is grown *in-vivo* on SP4 media. The *in-silico* media composition is based on the experimentally characterized composition, with additional essential molecules added (nucleobases, gases, polyamines, vitamins, and ions) in reported amounts to support *in-silico* cellular growth. Additionally, the *M. genitalium* whole-cell model represents 10 external stimuli including temperature, several types of radiation, and three stress conditions. For more information see Karr *et al*. Supplementary Tables S3F, S3H, S3R (Karr et al., 2012).

### Equipment

For the *M. genitalium* whole-cell model we used the University of Bristol Advanced Computing Research Centres’s BlueGem, a 900-core supercomputer, which uses the Slurm queuing system, to run whole-cell model simulations.

We used a standard office desktop computer, with 8GB of ram, to write new code, and interact with the supercomputer. We used the following GUI software on Windows 7: Notepad++ for code editing, Putty (ssh software) for terminal access to the supercomputer, FileZilla (ftp software) to move files in bulk to and from the supercomputer, and PyCharm (IDE software) as an inbuilt desktop terminal and for python debugging. The command line software used included: VIM for code editing, and SSH, Rsync, and Bash for communication and file transfer with the supercomputers.

### Data Format

For the *M. genitalium* whole-cell model the majority of output files are state-NNN.mat files, which are logs of the simulation split into 100-second segments. The data within a state-NNN.mat file is organised into the 16 cellular variables. These are typically arranged as 3-dimensional matrices or time series, which are flattened to conduct analysis. The other file types contain summaries of data spanning the simulation. Each gene manipulated simulation can consist of up to 500 files requiring between 0.4GB and 0.9GB. Each simulation takes 5 to 12 hours to complete in real time, 7 - 13.89 hours in simulated time.

### Data Analysis Process

For the *M. genitalium* whole-cell model, the raw data is automatically processed as the simulation ends. runGraphs.m carries out the initial analysis, while compareGraphs.m overlays the output on collated graphs of 200 unmodified *M. genitalium* simulations. Both outputs are saved as MATLAB .fig and .pdfs, though the .pdf files were the sole files analysed. The raw .mat files were stored in case of further investigation.

The GO biological process terms used for further analysis were downloaded from Uniprot (Apweiler et al., 2004) (strain ATCC 33530/NCTC 10195), processed by a created script (github.com/squishybinary/Gene_Ontology_Comparison_for_Mycoplasma_genitalium_whole-cell_model) in combination with lists of genes, organised manually into tables of GO terms that were unaffected, reduced, or removed entirely by gene deletions, and then analysed.

### Modelling Scripts

There are six scripts used to run the *M. genitalium* whole-cell model. Three are the experimental files created with each new experiment (the bash script, gene list, experiment list), and three are stored within the whole-cell model and are updated only upon improvement (MGGrunner.m, runGraphs.m, and compareGraphs.m). The bash script is a list of commands for the supercomputers) to carry out. Each bash script determines how many simulations to run, where to store the output, and where to store the results of the analysis. The gene list is a text file containing rows of gene codes (in the format ‘MG_XXX’,). Each row corresponds to a single simulation and determines which genes that simulation should knockout. The experiment list is a text file containing rows of simulation names. Each row corresponds to a single simulation and determines the final location of the simulation output and analysis results.

## Acknowledgements

We thank the Advanced Computing Research Centre (ACRC) and BrisSynBio, a BBSRC/EPSRC Synthetic Biology Research Centre, at the University of Bristol for access to the Bluegem supercomputer.

## Author Contributions

C.G., L.M., O.C., J.R-G were involved in ideation. O.C. conducted simulations comparing the Stanford and Rockville 2 sets, a prototype version of the research presented here. J.R-G. collated, simulated, analysed and repaired the nine sets, and wrote the paper and supplementary data 1 - 16, 26 - 29. J.R. analysed the nine sets and wrote supplementary data 17 - 25. C.G. and L.M supervised the project. C.G., L.M., J.R. edited the paper.

## References

Apweiler, R., Bairoch, A., Wu, C. H., Barker, W. C., Boeckmann, B., Ferro, S., Gasteiger, E., Huang, H., Lopez, R., Magrane, M., Martin, M. J., Natale, D. A., O’Donovan, C., Redaschi, N., & Yeh, L.-S. L. (2004). UniProt: the Universal Protein knowledgebase. Nucleic Acids Research, 32(Database issue), D115–D119.

Benders, G. A., Noskov, V. N., Denisova, E. A., Lartigue, C., Gibson, D. G., Assad-Garcia, N., Chuang, R. Y., Carrera, W., Moodie, M., Algire, M. A., Phan, Q., Alperovich, N., Vashee, S., Merryman, C., Venter, J. C., Smith, H. O., Glass, J. I., & Hutchison, C. A. (2010). Cloning whole bacterial genomes in yeast. Nucleic Acids Research, 38(8), 2558–2569.

du Plessis, D. J. F., Nouwen, N., & Driessen, A. J. M. (2011). The Sec translocase. Biochimica et Biophysica Acta, 1808(3), 851–865.

Dzieciol, A. J., & Mann, S. (2012). Designs for life: protocell models in the laboratory. Chemical Society Reviews, 41(1), 79–85.

Fleischmann, R. D., Adams, M. D., White, O., Clayton, R. A., Kirkness, E. F., Kerlavage, A. R., Bult, C. J., Tomb, J. F., Dougherty, B. A., & Merrick, J. M. (1995). Whole-genome random sequencing and assembly of Haemophilus influenzae Rd. Science, 269(5223), 496–512.

Forster, A. C., & Church, G. M. (2006). Towards synthesis of a minimal cell. Molecular Systems Biology, 2. https://doi.org/10.1038/msb4100090

Fraser, C. M., Gocayne, J. D., White, O., Adams, M. D., Clayton, R. A., Fleischmann, R. D., Bult, C. J., Kerlavage, A. R., Sutton, G., Kelley, J. M., Fritchman, J. L., Weidman, J. E., Small, K. V., Sandusky, M., Fuhrmann, J., Nguyen, D., Utterback, T. R., Saudek, D. M., Phillips, C. A. Venter, J. C. (1995). The Minimal Gene Complement of Mycoplasma-genitalium. Science, 270(5235), 397–403.

Fredens, J., Wang, K., de la Torre, D., Funke, L. F. H., Robertson, W. E., Christova, Y., Chia, T., Schmied, W. H., Dunkelmann, D. L., Beránek, V., Uttamapinant, C., Llamazares, A. G., Elliott, T. S., & Chin, J. W. (2019). Total synthesis of Escherichia coli with a recoded genome. Nature, 569(7757), 514–518.

Gibson, D. G., Benders, G. A., Andrews-Pfannkoch, C., Denisova, E. A., Baden-Tillson, H., Zaveri, J., Stockwell, T. B., Brownley, A., Thomas, D. W., Algire, M. A., Merryman, C., Young, L., Noskov, V. N., Glass, J. I., Venter, J. C., Hutchison, C. A., & Smith, H. O. (2008). Complete chemical synthesis, assembly, and cloning of a Mycoplasma genitalium genome. Science, 319(5867), 1215–1220.

Gibson, D. G., Benders, G. A., Axelrod, K. C., Zaveri, J., Algire, M. A., Moodie, M., Montague, M. G., Venter, J. C., Smith, H. O., & Hutchison, C. A. (2008). One-step assembly in yeast of 25 overlapping DNA fragments to form a complete synthetic Mycoplasma genitalium genome. Proceedings of the National Academy of Sciences of the United States of America, 105(51), 20404–20409.

Gibson, D. G., Glass, J. I., Lartigue, C., Noskov, V. N., Chuang, R. Y., Algire, M. A., Benders, G. A., Montague, M. G., Ma, L., Moodie, M. M., Merryman, C., Vashee, S., Krishnakumar, R., Assad-Garcia, N., Andrews-Pfannkoch, C., Denisova, E. A., Young, L., Qi, Z. Q., Segall-Shapiro, T. H. Venter, J. C. (2010). Creation of a Bacterial Cell Controlled by a Chemically Synthesized Genome. Science, 329(5987), 52–56.

Gil, R. (2015). The Minimal Gene-Set Machinery. In R. A. Meyers (Ed.), Synthetic Biology (pp. 443–478). John Wiley & Sons.

Gil, R., Silva, F. J., Pereto, J., & Moya, A. (2004). Determination of the core of a minimal bacterial gene set. Microbiology and Molecular Biology Reviews: MMBR, 68(3), 518–+.

Glass, J. I., Assad-Garcia, N., Alperovich, N., Yooseph, S., Lewis, M. R., Maruf, M., Hutchison, C. A., Smith, H. O., & Venter, J. C. (2006). Essential genes of a minimal bacterium. Proceedings of the National Academy of Sciences of the United States of America, 103(2), 425–430.

Glass, J. I., Merryman, C., Wise, K. S., Hutchison, C. A., 3rd, & Smith, H. O. (2017). Minimal Cells-Real and Imagined. Cold Spring Harbor Perspectives in Biology. https://doi.org/10.1101/cshperspect.a023861

Huang, C. H., Hsiang, T., & Trevors, J. T. (2013). Comparative bacterial genomics: defining the minimal core genome. Antonie Van Leeuwenhoek International Journal of General and Molecular Microbiology, 103(2), 385–398.

Hutchison, C. A., Chuang, R. Y., Noskov, V. N., Assad-Garcia, N., Deerinck, T. J., Ellisman, M. H., Gill, J., Kannan, K., Karas, B. J., Ma, L., Pelletier, J. F., Qi, Z. Q., Richter, R. A., Strychalski, E. A., Sun, L. J., Suzuki, Y., Tsvetanova, B., Wise, K. S., Smith, H. O., … Venter, J. C. (2016). Design and synthesis of a minimal bacterial genome. Science, 357(6280), 1414–U73.

Hutchison, C. A., Peterson, S. N., Gill, S. R., Cline, R. T., White, O., Fraser, C. M., Smith, H. O., & Venter, J. C. (1999). Global transposon mutagenesis and a minimal mycoplasma genome. Science, 286(5447), 2165–2169.

Jiang, W. Y., Bikard, D., Cox, D., Zhang, F., & Marraffini, L. A. (2013). RNA-guided editing of bacterial genomes using CRISPR-Cas systems. Nature Biotechnology, 37(3), 233–239.

Juhas, M., Eberl, L., & Glass, J. I. (2011). Essence of life: essential genes of minimal genomes. Trends in Cell Biology, 27(10), 562–568.

Kanehisa, M., & Goto, S. (2000). KEGG: kyoto encyclopedia of genes and genomes. Nucleic Acids Research, 28(1), 27–30.

Karas, B. J., Jablanovic, J., Sun, L. J., Ma, L., Goldgof, G. M., Stam, J., Ramon, A., Manary, M. J., Winzeler, E. A., Venter, J. C., Weyman, P. D., Gibson, D. G., Glass, J. I., Hutchison, C. A., Smith, H. O., & Suzuki, Y. (2013). Direct transfer of whole genomes from bacteria to yeast. Nature Methods, 10(5), 410–+.

Karr, J. R., Sanghvi, J. C., Macklin, D. N., Gutschow, M. V., Jacobs, J. M., Bolival, B., Jr, Assad-Garcia, N., Glass, J. I., & Covert, M. W. (2012). A whole-cell computational model predicts phenotype from genotype. Cell, 150(2), 389–401.

Kazakiewicz, D., Karr, J. R., Langner, K. M., & Plewczynski, D. (2015). A combined systems and structural modeling approach repositions antibiotics for Mycoplasma genitalium. Computational Biology and Chemistry, 59 Pt B, 91–97.

Koonin, E. V. (2003). Comparative genomics, minimal gene-sets and the last universal common ancestor. Nature Reviews. Microbiology, 1(2), 127–136.

Lagesen, K., Ussery, D. W., & Wassenaar, T. M. (2010). Genome update: the 1000th genome -a cautionary tale. Microbiology-Sgm, 156, 603–608.

Macklin, D. N., Ahn-Horst, T. A., Choi, H., Ruggero, N. A., Carrera, J., Mason, J. C., Sun, G., Agmon, E., DeFelice, M. M., Maayan, I., Lane, K., Spangler, R. K., Gillies, T. E., Pauli, M. L., Akhter, S., Bray, S. R., Weaver, D. S., Keseler, I. M., Karp, P. D., … Covert, M. W. (2020). Simultaneous cross-evaluation of heterogeneous E. coli datasets via mechanistic simulation. Science, 369(6502). https://doi.org/10.1126/science.aav3751

Mushegian, A. R., & Koonin, E. V. (1996). A minimal gene set for cellular life derived by comparison of complete bacterial genomes. Proceedings of the National Academy of Sciences of the United States of America, 93(19), 10268–10273.

Purcell, O., Jain, B., Karr, J. R., Covert, M. W., & Lu, T. K. (2013). Towards a whole-cell modeling approach for synthetic biology. Chaos, 23(2), 025112.

Rancati, G., Moffat, J., Typas, A., & Pavelka, N. (2018). Emerging and evolving concepts in gene essentiality. Nature Reviews. Genetics, 19(1), 34–49.

Rees-Garbutt, J., Chalkley, O., Landon, S., Purcell, O., Marucci, L., & Grierson, C. (2020). Designing minimal genomes using whole-cell models. Nature Communications, 11(1), 836.

Reich, K. A. (2000). The search for essential genes. Research in Microbiology, 151(5), 319–324.

Sanghvi, J. C., Regot, S., Carrasco, S., Karr, J. R., Gutschow, M. V., Bolival, B., Jr, & Covert, M. W. (2013). Accelerated discovery via a whole-cell model. Nature Methods, 10(12), 1192–1195.

Shuler, M. L., Foley, P., & Atlas, J. (2012). Modeling a Minimal Cell. In A. Navid (Ed.), Microbial Systems Biology (Vol. 881, pp. 573–610). Humana Press.

Tomita, M., Hashimoto, K., Takahashi, K., Shimizu, T. S., Matsuzaki, Y., Miyoshi, F., Saito, K., Tanida, S., Yugi, K., Venter, J. C., & Hutchison, C. A. (1999). E-CELL: software environment for whole-cell simulation. Bioinformatics, 15(1), 72–84.

Tsarmpopoulos, I., Gourgues, G., Blanchard, A., Vashee, S., Jores, J., Lartigue, C., & Sirand-Pugnet, P. (2016). In-Yeast Engineering of a Bacterial Genome Using CRISPR/Cas9. ACS Synthetic Biology, 5(1), 104–109.

